# PGC-1β maintains mitochondrial metabolism and restrains inflammatory gene expression

**DOI:** 10.1101/2022.05.18.492477

**Authors:** Hannah Guak, Ryan D. Sheldon, Ian Beddows, Alexandra Vander Ark, Hui Shen, Russell G. Jones, Julie St-Pierre, Eric H. Ma, Connie M. Krawczyk

## Abstract

Metabolic programming of the innate immune cells known as dendritic cells (DCs) changes in response to different stimuli, influencing their function. While the mechanisms behind increased glycolytic metabolism in response to inflammatory stimuli are well-studied, less is known about the programming of mitochondrial metabolism in DCs. We used lipopolysaccharide (LPS) and interferon-β (IFN-β), which differentially stimulate the use of glycolysis and oxidative phosphorylation (OXPHOS), respectively, to identify factors important for mitochondrial metabolism. We found that the expression of peroxisome proliferator-activated receptor gamma coactivator 1β (PGC-1β), a transcriptional co-activator and known regulator of mitochondrial metabolism, decreases when DCs are activated with LPS, when OXPHOS is diminished, but not with IFN-β, when OXPHOS is maintained. We examined the role of PGC-1β in bioenergetic metabolism of DCs and found that PGC-1β deficiency in DCs indeed impairs mitochondrial respiration. PGC-1β-deficient DCs are more glycolytic compared to controls, likely to compensate for reduced OXPHOS. PGC-1β deficiency also causes decreased capacity for ATP production at steady state and in response to IFN-β treatment. Loss of PGC-1β in DCs leads to increased expression of genes in inflammatory pathways, and reduced expression of genes encoding proteins important for mitochondrial metabolism and function. Collectively, these results demonstrate that PGC-1β is a key positive regulator of mitochondrial metabolism in DCs.

## Introduction

Numerous studies in recent years have established cellular metabolism as an essential regulator of immune cell function and activation ^1–3^. Two of the main ATP-generating pathways are glycolysis, which occurs in the cytosol, and oxidative phosphorylation (OXPHOS), which takes place in the mitochondria. During mitochondrial metabolism, the tricarboxylic (TCA) cycle produces reducing agents that fuel the electron transport chain (ETC) with oxygen as the final electron acceptor, thus consuming oxygen in the process of OXPHOS.

The peroxisome proliferator-activated receptor gamma coactivator 1 (PGC-1) group of transcriptional co-activators are major regulators of mitochondrial metabolism. This group comprises PGC-1α and PGC-1β, which are close homologs, and a less closely related member PGC-related coactivator (PRC) ^4^. PGC-1 proteins interact with and promote the activity of several transcription factors—including but not limited to PPARα/β/δ/γ, NRF1/2, ERRα/β/γ, and SREBP1a/1c/2—that regulate mitochondrial metabolism, lipid metabolism, and other aspects of cellular metabolism (reviewed in ^4^). PGC-1α and PGC-1β regulate the expression of overlapping sets of genes but can have distinct functions in different tissues. For example, in the liver, PGC-1α enhances the expression of genes for gluconeogenesis in response to fasting, while PGC-1β has a key role in lipogenesis and lipoprotein secretion in response to dietary fats ^5,6^. In addition, PGC-1β promotes cellular respiration that is more highly coupled to energy production in myoblasts compared to PGC-1α ^7^.

Existing studies on PGC-1 proteins in the context of immunity have largely been focused on PGC-1α, with a relatively small number on PGC-1β. For example, PGC-1α is important for promoting antitumor immunity by enforcing mitochondrial biosynthesis and metabolic fitness in T cells ^8,9^. In exhausted virus-specific T cells, overexpression of PGC-1α similarly helps reverse some of their metabolic defects and counteract exhaustion ^10^. PGC-1β is known, however, to promote oxidative metabolism to drive alternative activation (required for modulating inflammation and tissue repair) in response to IL-4 in macrophages ^11^.

Dendritic cells (DCs) are innate immune cells that are found in most tissues throughout the body, constantly patrolling and sampling their microenvironment. At steady state, DCs engage both glycolysis and oxidative metabolism ^1^. Upon Toll-like receptor (TLR) stimulation, DCs are known to rapidly upregulate their glycolytic activity, which is required for optimal DC activation ^12^. At late-stage activation, glycolysis maintains survival of TLR-activated DCs ^1^. Conversely, an increase in fatty acid oxidation (FAO) drives the generation of tolerogenic DCs ^13,14^. While the role of glycolysis in regulating DC function has been well studied ^1,12,15^, the role of mitochondrial metabolism in DCs requires more extensive investigation. Here, we characterized the metabolic and transcriptional changes that occur in DCs treated with lipopolysaccharide (LPS), bacterial component that induces glycolysis in DCs ^1,12^, or interferon-β (IFN-β), an antiviral mediator. We found that PGC-1β gene expression dramatically decreases with LPS stimulation, but levels are largely maintained in IFN-β-treated cells. PGC-1β deficiency impairs ATP production by oxidative metabolism and shifts the bioenergetic balance towards glycolysis. Gene expression analyses reveals that PGC-1β loss in DCs leads to an increase in inflammatory genes, and reduced expression of genes encoding proteins important for optimal mitochondrial function, including the TCA cycle, respiratory electron transport, and cristae formation. In all, we demonstrate the importance of PGC-1β in the maintenance of oxidative metabolism and bioenergetic capacity, and for limiting the activation state of DCs.

## Results

### Comparing metabolite and gene expression profiles of metabolically distinct DCs

To compare metabolite levels and gene expression in metabolically distinct cells, we used the bacterial component, LPS, and the antiviral mediator, IFN-β. We measured the extracellular acidification rate (ECAR) and oxygen consumption rate (OCR) of these cells using the Seahorse bioanalyzer, with sequential treatments with mitochondrial drugs to determine parameters of mitochondrial function. The results show that LPS- and IFN-β-treated DCs have contrasting bioenergetic profiles (Fig. 1a), with LPS-activated cells not using OXPHOS and therefore not responding to inhibitors and activators of mitochondrial respiration, as previously established ^16^ and IFN-β-treated DCs having greater maximal respiration than unstimulated DCs (Fig. 1a, left). ECAR for LPS-activated cells was greater than unstimulated cells, which is in line with previous results ^1,12^, while ECAR was not changed with IFN-β treatment (Fig. 1a, right).

**Figure 1.**
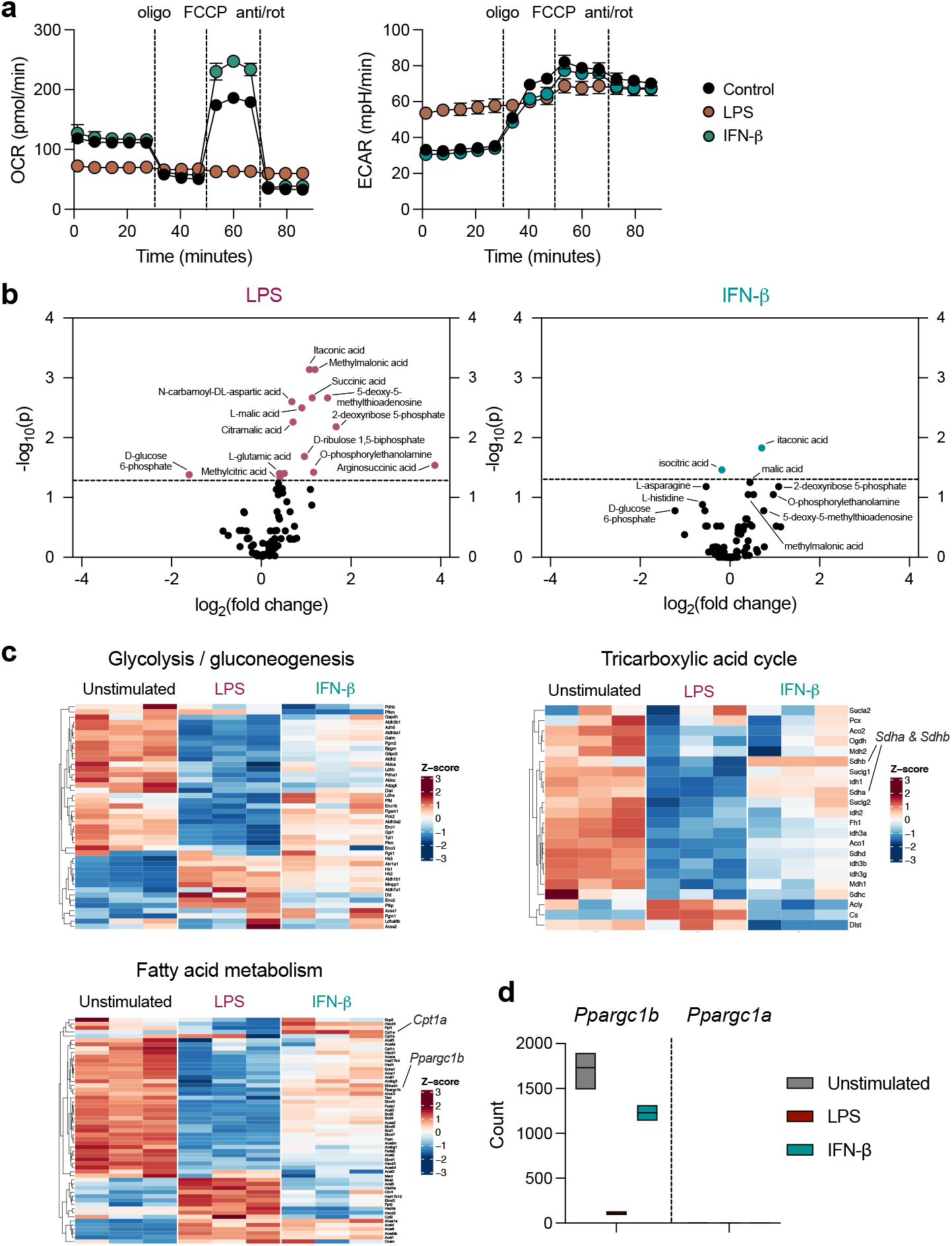
Comparing metabolite and gene expression profiles or metabolically distinct DCs. DCs were stimulated with LPS (100 ng/mL) or IFN-β (1000 U/mL) for 18 hours and the **(a)** OCR (left) and ECAR (right) were measured by Seahorse bioanalyzer over time with sequential treatments with oligomycin (oligo), FCCP, and antimycin and rotenone (anti/rot). **(b)** Volcano plot of data significance (-log10(p)), with a dotted line at *p* = 0.05, versus log_2_(fold change) for changes in metabolite relative abundance in LPS-activated DCs (left) and IFN-β-treated (right) compared to unstimulated DCs. **(c)** Heat maps of expression of genes related to glycolysis/gluconeogenesis, tricarboxylic acid cycle, and fatty acid metabolism, by unstimulated DCs, or DCs activated with LPS or IFN-β for 6 hours. **(d)** Raw counts from RNA sequencing data of *Ppargc1b* and *Ppargc1a*. Data in **(a)** are of one experiment representative of at least three experiments (mean and s.d. of five replicates).

We compared the metabolite profiles of cells treated with LPS or IFN-β using LC-MS (Fig. 1b) and analysis of genes encoding factors involved in metabolic pathways (Fig. 1c). LPS induced significant changes in metabolites involved in glycolysis, the TCA cycle, pentose phosphate pathway, and amino acid metabolism, while changes induced by IFN-β were minimal (Fig 1b). Despite their contrasting OCR, ECAR, and metabolite profiles, similar genes encoding proteins involved in major metabolic pathways, including glycolysis, the TCA cycle, and fatty acid metabolism, showed changes in expression in response to LPS or IFN-β treatment (Fig 1c). However, the overall magnitude of change for a given gene induced by IFN-β was less. Some notable exceptions include the maintenance of *Sdha* and *Sdhb* expression, which are members of complex II of the ETC, and the induction of *Cpt1a* expression by IFN-β treatment, which is consistent with previous studies ^17,18^. *Ppargc1b*, which encodes PGC-1β, a transcriptional co-activator important for mitochondrial metabolism, was also maintained in IFN-β-treated DCs but decreased in LPS-stimulated DCs. *Ppargc1a* was not detected at baseline, nor was its expression induced by either LPS or IFN-β treatment (Fig. 1d). We validated the RNA sequencing results using RT-qPCR and confirmed that *Ppargc1b* is downregulated by LPS but maintained during IFN-β treatment compared with untreated cells (Supplementary Fig. S1).

### PGC-1β-deficient DCs have impaired oxidative metabolism and increased glycolysis

To determine how PGC-1β affects mitochondrial metabolism, we first examined bioenergetic metabolism in DCs, with cells expressing an shRNA targeting *Ppargc1b* (*shPpargc1b;* Supplementary Fig. S2). Knockdown efficiency of two different shRNA for *Ppargc1b* were determined by RT-qPCR (Supplementary Fig. S2a,b). *shPpargc1b* #1 is used for most experiments presented unless stated otherwise. DCs transduced with sh*Ppargc1b* have reduced basal OCR and spare respiratory capacity compared to the control (SRC; difference between maximal OCR and basal OCR) (Fig. 2a; Supplementary Fig. S2c). In contrast, the basal ECAR of PGC-1β-deficient DCs is higher (Fig. 2b). FCCP treatment further elevates ECAR, but fails to substantially raise OCR in PGC-1β-deficient DCs (Fig. 2b), suggesting that at basal rates, these cells are already operating at maximal oxidative capacity.

**Figure 2.**
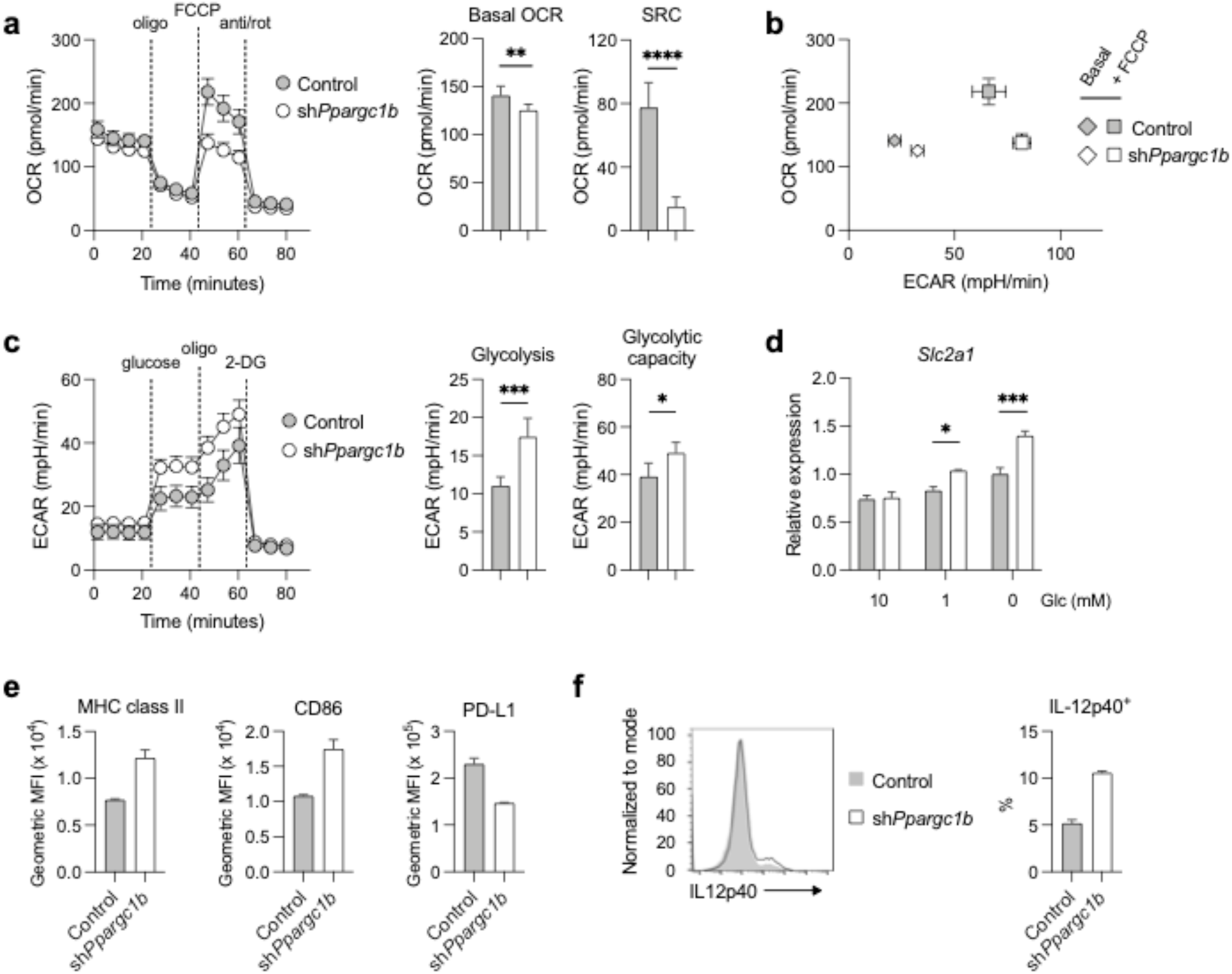
PGC-1β-deficient DCs have impaired oxidative metabolism and increased glycolysis. DCs were transduced with control shRNA or *Ppargc1b* shRNA and **(a)** OCR was measured by Seahorse bioanalyzer over time with sequential treatments with oligomycin (oligo), FCCP, and antimycin and rotenone (anti/rot). Basal oxygen consumption rate (OCR) and spare respiratory capacity (SRC) are determined from these measurements (right). **(b)** Basal and maximal (with FCCP) levels of OCR and ECAR from experiment in **(a)**. **(c)** ECAR was measured over time with sequential treatments with glucose, oligomycin, and 2-deoxyglucose (2-DG). From this graph (left), glycolysis (difference between ECAR after glucose addition and non-glycolytic ECAR before glucose addition) and glycolytic capacity (maximal ECAR after oligomycin treatment) were determined. **(d)** Gene expression of GLUT1 gene *Slc2a1* relative to *Hprt* levels in DCs cultured in 10, 1, or 0 mM glucose for 6 hours. **(e)** Geometric MFI of MHC class II, CD86, and PD-L1, and **(f)** intracellular levels of cytokine IL-12p40 (histogram on left; IL-12p40^+^ cells expressed as a percentage of transduced DCs on right) were determine by flow cytometry. Data in **(a-c)** are of one experiment representative of **(a-b)** five experiments, **(c)** three experiments (mean and s.d. of five to six replicates per condition), and **(e,f)** three experiments (mean and s.d. of duplicates). Data in **(d)** are of one experiment representative of two experiments (mean and s.e.m. of triplicates). Statistical significance was determined by **(a,c)** unpaired t-test or **(d)** two-way ANOVA. * p < 0.05, ** p < 0.01, *** p < 0.001, **** p < 0.0001

The increase in ECAR in PGC-1β-deficient DCs is likely due to increased glycolytic activity to compensate for the reduction in oxidative metabolism. To test the glycolytic capacity of DCs, we used sequential addition of glucose, oligomycin, and 2-deoxyglucose (2-DG) using the Seahorse bioanalyzer (Fig. 2c). Non-glycolytic ECAR (after 2-DG addition) was not different following PGC-1β deletion, but ECAR was higher when glucose was added (glycolysis; Fig. 2c). We also found that PGC-1β-deficient DCs have increased glycolytic capacity, measured by the maximal ECAR after addition of oligomycin, which inhibits mitochondrial respiration. This suggests that PGC-1β-deficient cells are programmed to use glycolysis for bioenergetic metabolism (Fig. 2c). In addition, gene expression of glucose transporter 1 (*Slc2a1*) is increased in PGC-1β-deficient DCs compared to control DCs under conditions of glucose withdrawal (Fig. 2d), suggesting increased dependence by the cells on glucose. Together, these results demonstrate that deletion of PGC-1β results in decreased mitochondrial oxidative capacity, and therefore a shift to glycolytic metabolism.

Since elevated glycolysis is required for optimal DC activation ^12,19^, we next examined whether PGC-1β-deficient DCs display enhanced activation levels. First, we determined that PGC-1β-deficient DCs express higher levels of MHC class II, a molecule for antigen presentation, and CD86, a costimulatory molecule, and reduced levels of PD-L1, an immunomodulatory ligand, compared to control DCs (Fig. 2e). In addition, at steady state, PGC-1β-deficient DCs express higher levels of the pro-inflammatory cytokines IL-12p40 (Fig. 2f). Thus, PGC-1β deficiency leads to higher baseline activation of DCs.

### PGC-1β-deficient DCs have reduced ATP production by oxidative metabolism

Given the shifts in bioenergetic preferences we observed, we next sought to understand whether PGC-1β deficiency impacts global cellular ATP production. ATP production rates attributed to oxidative phosphorylation and glycolysis were calculated using Seahorse assay measurements (see Methods). As expected, the basal ATP production rate from OXPHOS is reduced in PGC-1β-deficient cells, whereas ATP production from glycolysis is increased (Fig. 3a), demonstrating that in the absence of PGC-1β, cells can pivot to use glycolysis to support their bioenergetic needs. At maximal oxidative capacity (with FCCP treatment), the estimated ATP production by OXPHOS decreased in PGC-1β-deficient cells (Fig. 3a, right). However, compensation by glycolysis is not observed, resulting in overall decreased ATP production rates (Fig. 3b). These results demonstrate that while PGC-1β-deficient cells can engage glycolysis to compensate for loss of OXPHOS-generated ATP, this compensation is not complete.

**Figure 3.**
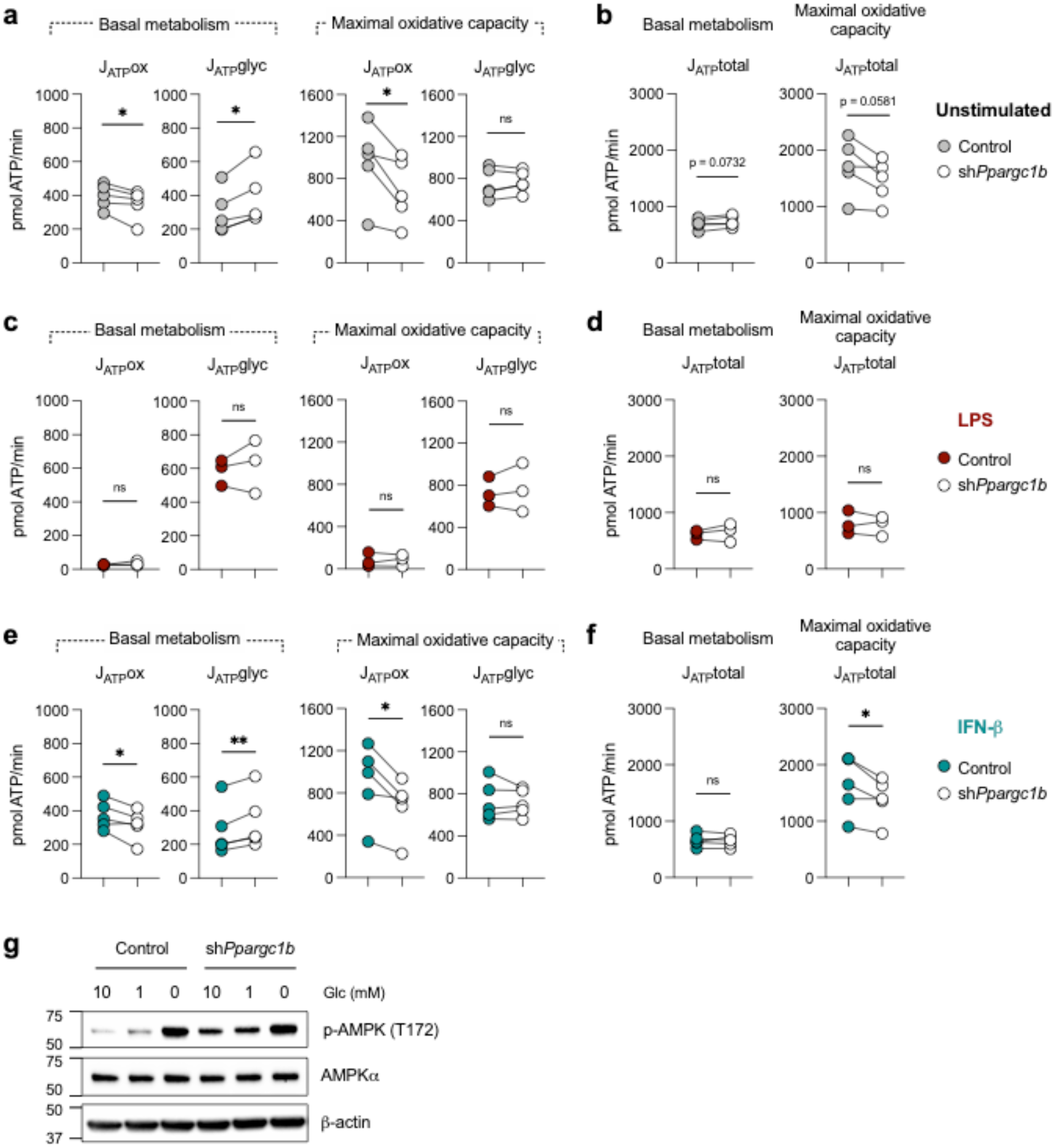
Effect of PGC-1β on metabolic phenotype of differentially-activated DCs. DCs were transduced with control shRNA or *Ppargc1b* shRNA and **(a)** ATP production rates by oxidative metabolism (J_ATP_ox) and glycolysis (J_ATP_glyc) were calculated from Seahorse assay measurements (see Methods) at basal (left) and maximal (right) respiratory rates. **(b)** Total ATP production rates (J_ATP_total) at basal metabolism and maximal respiration were determined by adding J_ATP_ox and J_ATP_glyc from **(a)**. ATP production rates by oxidative metabolism (J_ATP_ox) and glycolysis (J_ATP_glyc) were calculated from Seahorse assay measurements in DCs stimulated with **(c,d)** LPS (100 ng/mL) or **(e,f)** IFN-β (1000 U/mL). Total ATP production rates of stimulated DCs in **(d,f)** at basal metabolism and maximal respiration were determined by adding J_ATP_ox and J_ATP_glyc from **(c,e)**. Colored circles are control DCs and open circles are DCs transduced with *Ppargc1b* shRNA. **(g)** AMPK phosphorylated at threonine 172 (p-AMPK T172), AMPKa, and β-actin, were visualized by western blot for DCs that were cultured in media containing 10, 1, or 0 mM glucose. Data are of **(c,d)** three or **(a,b,e,f)** five experiments pooled together, with each individual experiment represented by a circle and values from the same experiment joined by a line. Data in **(g)** are of one experiment representative of three experiments. Statistical significance was determined by paired *t*-test. * p < 0.05, ** p < 0.01

We next investigated how PGC-1β deficiency affects the metabolic phenotypes of activated DCs. Since stimulation by LPS results in downregulation of *Ppargc1b* expression (Fig. 1d), we hypothesized that PGC-1β deficiency would not have a considerable effect on the metabolism of LPS-activated DCs. IFN-β-treated DCs, in contrast, maintain PGC-1β expression, and therefore were predicted to be affected by PGC-1β deficiency. Both basal and maximal ATP production rates by LPS-activated DCs were not significantly affected by PGC-1β deficiency (Fig. 3c). Total ATP production rates by LPS-activated DCs were also not significantly different (Fig. 3d). ATP production rates by oxidative metabolism in IFN-β-treated DCs lacking PGC-1β, however, were lower at both basal metabolism and maximal respiration (Fig. 3e). Basal ATP production rates by glycolysis were higher in PGC-1β-deficient DCs treated with IFN-β (Fig. 3e). Like that of unstimulated DCs, total ATP production rates at maximal oxidative capacity by IFN-β-treated DCs were lower in PGC-1β-deficient DCs (Fig. 3f). These data confirm that PGC-1β is required for optimal bioenergetic output in activated DCs that use oxidative metabolism.

Disruption of energy homeostasis can activate the energy sensor 5’ adenosine monophosphate-activated protein kinase (AMPK), which in turn activates metabolic pathways such as glycolysis and OXPHOS to promote the maintenance of ATP levels ^20,21^. Since the ATP production rate by oxidative metabolism is blunted by PGC-1β loss, we examined AMPK activation by measuring the levels of phosphorylation at residue threonine-172 (T172). We confirmed that phospho-AMPK at T172 increases when glucose is restricted (Fig. 3g). We found that PGC-1β-deficient DCs have higher levels of phosphorylation at T172 compared to control DCs (Fig. 3g), indicating that the cells sense diminished ATP and AMPK is activated to counteract the energy loss resulting from PGC-1β deficiency. Despite the energy stress, we confirmed that PGC-1β deficiency does not lead to a significant change in cell death at steady state (Supplementary Fig. S3a).

AMPK is also known to promote autophagy, which can make available more nutrients for cells to catabolize ^22^; we therefore measured markers of autophagy, sequestosome 1 (SQSTM1 or p62) and microtubule-associated protein 1 light chain 3 (LC3). Cells were cultured in either full glucose conditions or in the absence of glucose to induce nutrient stress. Chloroquine treatment was used to determine the flux of autophagy, as it blocks autophagosome-lysosome fusion, resulting in accumulation of autophagosome proteins. The expression of autophagy-related proteins SQSTM1 and LC3 was elevated in PGC-1β-deficient DCs (Supplementary Fig. S3b,d). Gene expression of SQSTM1 is also greater with glucose deprivation (Supplementary Fig. S3c). LC3 is involved in autophagosome formation, with LC3-I as the cytoplasmic form and LC3-II as the modified form that associates with autophagosomes. The relative intensity of LC3-II normalized to β-actin levels shows that LC3-II levels are greater in PGC-1β-deficient cells (Supplementary Fig. S3d). Autophagic flux, measured by determining the difference in intensity of LC3-II in the presence or absence of chloroquine for a given condition, is elevated in PGC-1β-deficient cells as well (Supplementary Fig. S3d). This suggests that the metabolic stress resulting from PGC-1β deficiency also stimulates autophagy as cells try to maintain energy homeostasis.

### Impaired oxidative metabolism in PGC-1β-deficient DCs is not due to net changes in mitochondrial mass nor is it substrate-specific

Since PGC-1 co-activators are known to be important for mitochondrial biogenesis in many tissues ^23^, and we have shown that autophagy increases with PGC-1β deficiency (Supplementary Fig. S3b-d), we examined whether reduced mitochondrial content is responsible for the impaired OCR by PGC-1β-deficient DCs. By flow cytometry, we determined that the mean fluorescence intensity (MFI) of dyes to measure mitochondrial mass is not reduced in PGC-1β-deficient DCs (Fig. 4a). The levels of voltage-dependent anion channel (VDAC), typically used as a loading control for mitochondrial content, was also unchanged (Fig. 4b). Despite PGC-1β-deficient DCs having reduced gene expression of many TCA cycle and ETC-related proteins, protein levels of the ETC complexes were not noticeably different (Fig. 4b) suggesting there is a functional defect in the mitochondria.

**Figure 4.**
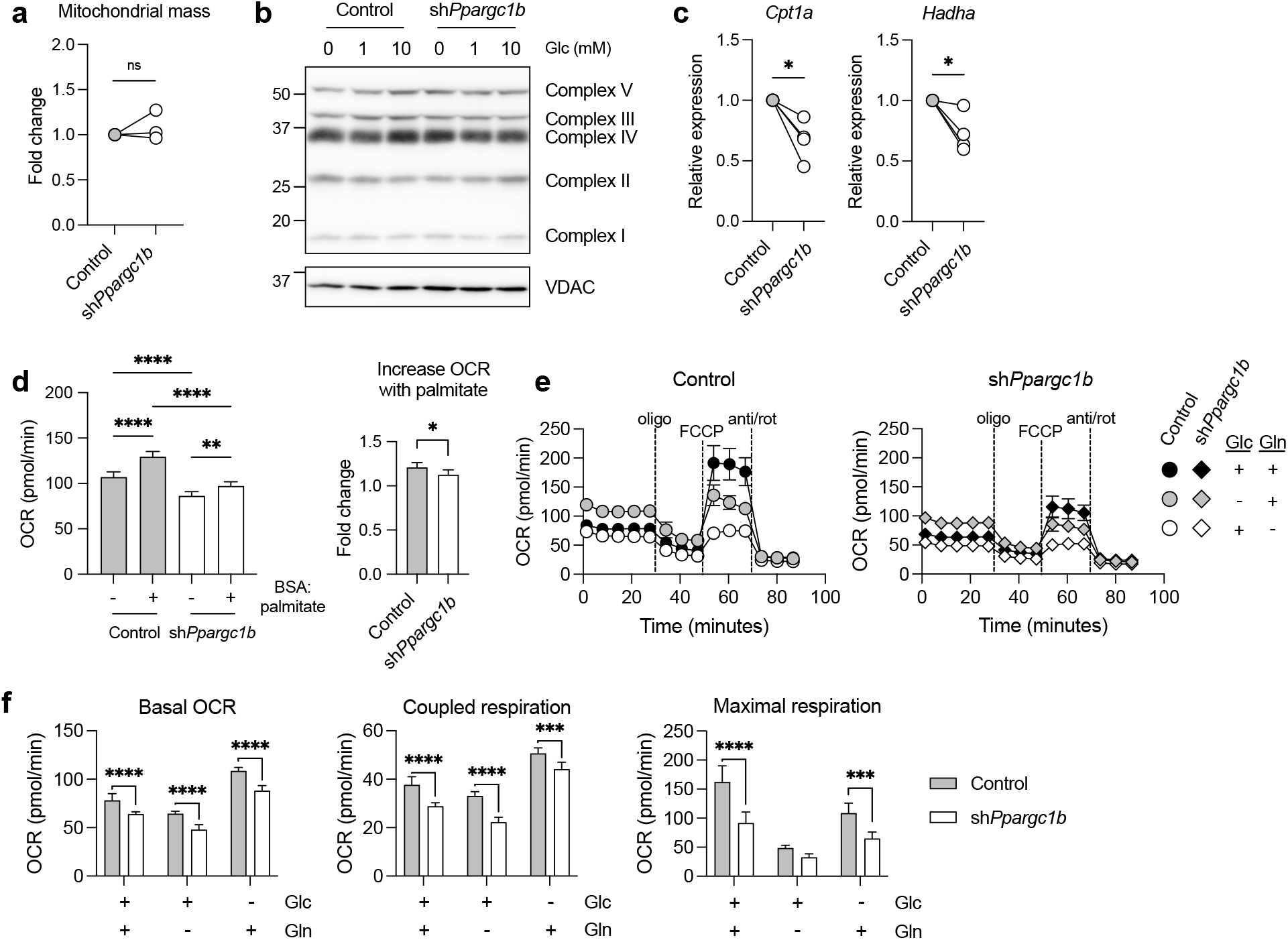
PGC-1β-deficient DCs exhibit reduced fatty acid oxidation, but unchanged total mitochondrial mass. DCs were transduced with control shRNA or *Ppargc1b* shRNA. **(a)** Mitochondrial mass of DCs stained with MitoSpy Green or MitoTracker Deep Red, with geometric MFIs represented as fold changes relative to the control condition. **(b)** Protein expression of complexes I-V of the ETC and VDAC of control or PGC-1β-deficient DCs cultured with 0, 1, or 10 mM glucose for 6 hours. **(c)** Gene expression fold changes of *Cpt1a* (left) and *Hadha* (right) relative to *Hprt*. **(d)** DCs were cultured in nutrient-limiting media for 6 hours and OCR was measured immediately following addition of either BSA or BSA-conjugated palmitate. **(e)** OCR was measured by Seahorse bioanalyzer over time with sequential treatments with oligomycin (oligo), FCCP, and antimycin and rotenone (anti/rot), in assay medium with or without 10 mM glucose or 2 mM glutamine. **(f)** Basal OCR, coupled respiration, and maximal respiration determined from the experiment in **(e)**. Data from **(a,c)** are of **(a)** three and **(c)** four experiments pooled together, with each individual experiment represented by a circle. Data from **(b)** is one experiment representative of three experiments. Data from **(d-f)** are of one experiment representative of two experiments (s.d. of five to six replicates per condition). Statistical significance was determined by **(a,c)** paired *t*-test, **(d, left)** one-way ANOVA, **(d, right)** unpaired *t*-test, or **(e,g)** two-way ANOVA. * p < 0.05, ** p < 0.01, *** p < 0.001, **** p < 0.0001

Processing of different substrates, including fatty acids, can contribute to the OCR. Since PGC-1β is known to target transcription factors that regulate lipid metabolism ^6,24^, we examined whether components involved in FAO were affected by PGC-1β deficiency. We determined that gene expression of *Hadha* and *Cpt1a*, which encode enzymes involved in key steps of FAO, were reduced in PGC-1β-deficient DCs (Fig. 4c). In addition, control DCs more readily utilize fatty acids as fuel after nutrient starvation compared to PGC-1β-deficient DCs, as determined by OCR when the cells are in the presence of BSA-conjugated palmitate (Fig. 4c). Together, these data suggest that the reduced oxidative metabolism by PGC-1β-deficient DCs is in part due to impaired FAO, and not due to diminished mitochondrial biosynthesis.

We also assessed how nutrient limitation of other carbon sources affects OCR. In control cells, removing glucose results in an increase in basal OCR (likely to compensate for the loss of ATP production by glycolysis), but a decrease in maximal respiration, whereas removal of glutamine results in a decrease in both basal OCR and maximal respiration (Fig. 4e). These data demonstrate that glutamine is required for optimal maximal respiration even in the presence or absence of glucose. For all conditions, PGC-1β-deficient cells have lower basal and maximal respiration compared to control DCs (Fig. 4e,f). Altogether, these data indicate that PGC-1β-deficient cells have reduced oxidative metabolism compared to the control regardless of the substrate, suggesting that the mitochondria-specific dysfunction is substrate-independent.

### PGC-1β loss results in succinate accumulation

While PGC-1β-deficient DCs have increased glycolysis (Fig. 2c), they also have reduced glucose-supported maximal respiration (Fig. 4e,f). We therefore examined whether there is a difference in the proportions of glucose-derived TCA cycle metabolites by stable isotope labeling and metabolomics using ^13^C-glucose. We found that the proportion of glucose-derived TCA cycle metabolites is smaller in PGC-1β-deficient DCs (Fig. 5b), which is in line with our data showing less oxidation of glucose in these cells (Fig. 4e). Furthermore, the relative abundance of lactate (m+3 isotopologue) is elevated in PGC-1β-deficient DCs (Supplementary Fig. S4a), supporting the earlier results in Fig. 2 that these cells are more glycolytic. We also observed that the relative abundance of succinate is significantly elevated in the PGC-1β-deficient DCs (Fig. 5c). Relative abundances of several other TCA intermediates, including citrate, aconitate, and malate, also trend higher, while levels of α-ketoglutarate (α-KG) are unchanged (Fig. 5c). The ratio of succinate to malate (downstream of succinate in the TCA cycle) is higher in PGC-1β-deficient DCs (Fig. 5d), and the ratio of α-KG (upstream of succinate in the TCA cycle) to succinate is lower (Fig. 5e), demonstrating that there is an accumulation of succinate in PGC-1β-deficient cells. We examined the mitochondria by electron microscopy and found that mitochondria in PGC-1β-deficient DCs are less electron dense (Fig. 5f), suggesting reduced density of electron transport chain components. Because we observed normal protein levels of ETC components by western blot (Fig. 4b), we performed RNA-seq to determine the expression levels of other components of the TCA cycle and ETC. We found that at steady state and in response to IFN-β treatment, PGC-1β deficiency results in less expression of genes important for the TCA cycle and respiratory electron transport (Fig. 5g,h; S4b). PGC-1β-deficient DCs also exhibit an increase in gene expression in several inflammatory pathways (Fig. 5g; S4c), consistent with their increased activation phenotype (Fig 2e,f). Interestingly, genes important for cristae formation were also decreased in the absence of PGC-1β (Fig. 5h). One of the affected genes, *Chchd6*, has been shown to be critical for preserving the structural integrity of mitochondrial cristae, as its loss induces mitochondrial dysfunction ^25^. Together, these data demonstrate that PGC-1β is necessary for optimal oxidative metabolism and a functional ETC, which restrains the inflammatory phenotype of DCs.

**Figure 5.**
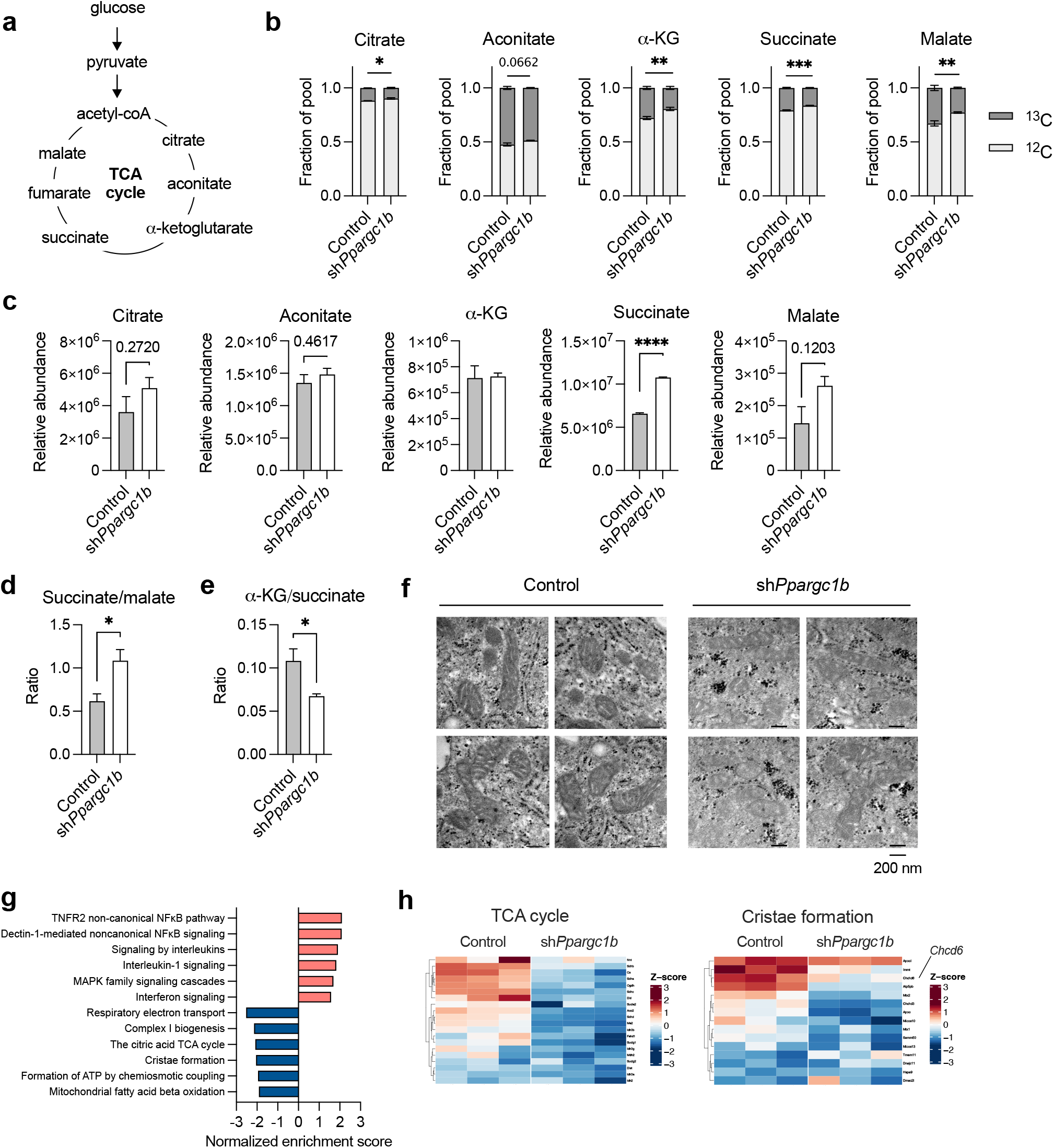
Succinate accumulates in PGC-1β-deficient DCs. **(a)** Schematic of glucose metabolism and TCA cycle. DCs were transduced with control shRNA or *Ppargc1b* shRNA and **(b)** fractional enrichment of select glucose-derived TCA cycle metabolites was measured following culture with U-[^13^C]-glucose for 6 hours. **(c)** Relative abundance of metabolites were measured by LC-MS (aconitate, α-KG, and succinate) or GC-MS (citrate and malate). **(d)** Ratio of succinate to malate from measurements in **(c)**. **(e)** Ratio of α-KG to succinate from measurements in **(c)**. **(f)** Mitochondria imaged by transmission electron microscopy (dark dots are microbeads used for sorting). **(g)** Select inflammatory and metabolic Reactome pathways that are significantly enriched in PGC-1β-deficient versus control DCs. **(h)** Heat maps of the expression of genes from the TCA cycle (left) and cristae formation (right) Reactome pathways. Data from **(b-e)** are of one experiment representative of two experiments (s.e.m. of three replicates). Statistical analysis in **(b)** was determined by 2-way ANOVA and represents the comparison of the ^13^C-labeled fraction between each condition. Statistical significance for **(c-e)** was determined by unpaired *t*-test. * p < 0.05, ** p < 0.01, *** p < 0.001, **** p < 0.0001

## Discussion

Previous studies have demonstrated that cellular metabolism is crucial for the regulation of immune cell function and activation ^2,3^. In this study, we found that in bone marrow-derived DCs, *Ppargc1a* is not appreciably expressed while *Ppargc1b* expression is dynamically changed in response to stimulation. We showed that LPS, but not IFN-β, induces downregulation of *Ppargc1b* expression, which we found notable, since in certain tissues such as skeletal muscle, PGC-1α expression is typically more dynamic, while PGC-1β expression is largely unchanged ^26,27^. PGC-1β deletion resulted in higher baseline activation in DCs, which supports previous studies that associate PGC-1β expression with an anti-inflammatory state ^28–30^.

We found that loss of PGC-1β expression leads to impaired oxidative metabolism and increases glycolytic activity to compensate. PGC-1β is necessary to maintain ATP levels in IFN-β-treated DCs, but not the ATP levels of LPS-activated DCs, which solely depend on glycolysis. In response to this energy deficiency in PGC-1β-deficient DCs, AMPK is activated (Fig. 3g), which likely leads to the upregulation of glycolysis and glucose uptake. Therefore, the increase in glycolytic rates in these cells is a compensatory mechanism to maintain intracellular ATP levels in response to the drop in ATP production by the mitochondria. While gene expression of glucose transporter *Slc2a1* in PGC-1β-deficient DCs is comparable to that of control DCs in full glucose conditions, glucose deprivation results in a greater fold change in *Slc2a1* expression, suggesting an increased reliance on glucose. These results highlight the impaired oxidative metabolism of PGC-1β-deficient DCs and their increased reliance on glycolysis.

Stress signals due to bioenergetic insufficiencies in PGC-1β-deficient DCs lead to AMPK activation, which is known to promote autophagy ^22^. Our data show that autophagy is indeed elevated in PGC-1β-deficient DCs (Supplementary Fig. S3). This result is in contrast with a few studies showing positive regulation of autophagy by PGC-1α and PGC-1β in muscle cells: PGC-1α deficiency results in a reduction in autophagy, and PGC-1β overexpression upregulates autophagy-related genes ^31,32^. Similarly, another study describes that the upregulation of autophagy-related genes induced in skeletal muscle cells by exercise is attenuated in PGC-1α-deficient mice ^33^. The most likely explanation of these differences is that in our study, autophagy is induced by AMPK activation and this is not due to specific transcriptional effects of PGC-1β loss.

To determine how PGC-1β deficiency leads to impaired oxidative metabolism, we first assessed mitochondrial content since PGC-1 proteins are important for mitochondrial biogenesis in certain contexts. Considering the elevated autophagy, we examined whether this increased autophagy leads to reduced mitochondria in PGC-1β-deficient DCs. However, mitochondrial mass was not consistently different between control and PGC-1β-deficient DCs, nor were the levels of select ETC components (Fig. 4a,b). We found that multiple substrates, including fatty acids, glucose, and glutamine, contribute to supporting oxidative metabolism in DCs (Fig. 4d-f). PGC-1β-deficient DCs, however, have reduced capacity to use each of these substrates for oxidative metabolism, suggesting there is a mitochondria-specific defect at the ETC or TCA cycle. Metabolite analysis shows a specific accumulation of succinate (Fig. 5), which indicates that the activity of succinate dehydrogenase, which is part of both the ETC and the TCA cycle, may also be defective. In addition, images of mitochondria acquired by electron microscopy reveal that mitochondria from PGC-1β-deficient DCs are less electron dense than those from control DCs (Fig. 5f), suggesting there is reduced electron flow through the ETC. Gene expression analyses reveal that several components of the ETC and the TCA cycle, and genes involved in cristae formation, show decreased expression. Together these data suggest that PGC-1β-deficient DCs have a dysfunctional cristae and ETC, leading to general impairment of mitochondrial metabolism.

The ETC has crucial roles in the inflammatory phenotype or differentiation of many immune cell types. For example, macrophages undergo remodeling of their ETC upon LPS activation that disrupts OXPHOS. This leads to production of reactive oxygen species (ROS) mainly by complex I and III, and the consequent stabilization of the transcription factor HIF-1a and expression of proinflammatory gene targets such as IL-1β and genes related to glycolysis ^34^. For T cells, complex II activity is required for optimal T helper (Th) 1 cell differentiation and IFN-γ production, while also suppressing their proliferation ^35^. ATP synthase (complex V) activity is necessary for Th17 differentiation and function, and inhibition of ATP synthase shifted CD4^+^ T cell differentiation into suppressive regulatory T cells ^36^. These and other studies illustrate that there are specific requirements of ETC complexes in immune cell function.

Since we found that the protein expression levels of several ETC components are not affected by PGC-1β loss in DCs, other mechanisms may be responsible for the dysfunctional ETC. Each of the ETC complexes are comprised of several protein subunits and protein expression of only select subunits were examined in this study. Optimal enzymatic activity of the ETC complexes requires precise assembly of their components, which involves not only proper expression of the subunits, but also a myriad of nuclear-encoded assembly factors ^37^. These assembly factors have a range of functions, including stabilization of the complexes and incorporation of necessary co-factors ^37^, several of which were expressed lower in PGC-1β -deficiency. Larger structures of ETC complexes known as supercomplexes enhance electron flux ^38^ and defective assembly of supercomplexes could provide an explanation for the reduced oxidative metabolism yet unchanged protein levels of individual ETC complexes. Further, reducing agents such as NADH and FADH_2_ are generated by the TCA cycle and donate electrons to the ETC, fueling OXPHOS. Diminished levels of these electron donors could result in reduced electron flow through the ETC. In addition, since gene expression for several proteins involved in cristae formation was decreased in PGC-1β deficiency (Fig. 5h), efficient organization and packing of the ETC components may also be affected. However, the precise mechanism by which PGC-1β supports efficient ETC activity in DCs requires additional investigation. Further studies to address these questions will help expand our findings that highlight the importance of PGC-1β in maintaining an oxidative metabolic program and bioenergetic balance in DCs.

## Methods

### Mice

C57BL/6 were bred and maintained at the Van Andel Research Institute in a specific pathogen-free environment. *Nos2-/-* mice were obtained from Jackson Laboratory. Up to four mice per cage were housed in microisolator cages on 12 hour light/dark cycles in a temperature-controlled environment with enviro-dry enrichment and free access to water and standard chow. Mice were subject to regular cage changes, veterinary inspection, and sentinel monitoring. Female mice were used at 8 to 20 weeks of age. All animal studies were carried out under protocols approved by the Van Andel Research Institute Institutional Animal Care and Use Committee and in accordance with ARRIVE guidelines.

### In vitro differentiation and retroviral transduction of dendritic cells

Bone marrow from the tibia and femur of C57BL/6 mice was extracted and seeded at day 0 in 6-well non-tissue culture-treated plates at 7.5 x 10^5^ non-erythrocytes/well or at 1 x 10^6^ non-erythrocytes/well if the cells were to undergo retroviral transduction. The cells were cultured in the presence of 20 ng/mL of granulocyte-macrophage colony-stimulating factor (GM-CSF; from Peprotech) in complete DC medium (CDCM): RPMI-1640 medium (Corning) containing 100 U/mL penicillinstreptomycin (Gibco), 2 mM L-glutamine (Gibco), 55 μM β-mercaptoethanol (Gibco), and 10% heat-inactivated Nu-Serum (Corning), with media changes at days 3 and 6. For retroviral transductions, the sequences for the *Ppargc1b* short hairpin RNA (shRNA) were obtained from RNAi Codex and cloned into the LMP retroviral vector expressing a human CD8 reporter ^39^. Retrovirus containing a control shRNA (firefly luciferase) or the *Ppargc1b* shRNA were produced by 293T cells transfected using Lipofectamine 2000 (Invitrogen). After 48 hours, supernatant containing retrovirus was collected and applied to day 2 DC culture and spun at 2500 rpm 30°C for 90 minutes. At day 8 or 9, DCs were harvested and sorted on human CD8 (for retrovirally transduced DCs) using anti-biotin microbeads (Miltenyi). Sorted cells were seeded at a final density of 1 x 10^6^ cells/mL and were left unstimulated or stimulated with 100 ng/mL LPS (*Escherichia coli* O111:B4; Invivogen) or 1000 U/mL IFN-β (PBL Interferon Source) for the indicated length of time.

### Transmission electron microscopy

Bone marrow-derived dendritic cells retrovirally transduced with control shRNA or *Ppargc1b* shRNA were sorted on human CD8 as described above. Cells were fixed in 2.5% glutaraldehyde in PBS and shipped to the Center for Advanced Microscopy at Michigan State University for analysis.

### Metabolic assay

DCs were analyzed by Seahorse XFe96 Analyzer (Agilent) to measure ECAR and OCR in realtime. DCs were seeded at 7 x 10^4^ cells/well in CDCM and stimulated for the indicated length of time. For the XF Cell Mito Stress Test (Agilent), the cell medium was replaced with XF RPMI base medium containing 10 mM glucose, 2 mM glutamine, and 5% FBS, with pH adjusted to 7.4. The DCs were sequentially treated with oligomycin (1.5 μM), FCCP (1.5 μM), antimycin/rotenone (both 1 μM), and monensin (20 μM). For the XF Glycolysis Stress Test (Agilent), the cell medium was replaced with XF RPMI base medium containing 2 mM glutamine and 5% FBS, but no glucose. The DCs were sequentially treated with glucose (10 mM), oligomycin (1.5 μM), and 2-deoxyglucose (50 mM). For the XF Palmitate Oxidation Stress Test (Agilent), DCs were cultured for 6 hours in nutrient-limiting media (RPMI-1640 containing 0.5 mM glucose, 1 mM L-glutamine, 1% FBS, 55 μM β-mercaptoethanol, 100 U/mL penicillin-streptomycin, and 0.5 mM carnitine), followed by replacing this with XF RPMI base medium containing 2 mM glucose and 0.5 mM carnitine. BSA or BSA-conjugated palmitate was added immediately prior to starting the run. For all assays, cells were placed at 37°C in a non-CO_2_ incubator for approximately 1 hour prior to the start of each run. After each run, cells were stained with 20 μM of Hoescht stain (ThermoFisher Scientific) for 15 minutes at 37°C and imaged using a Cytation imaging reader (BioTek). Measurements from the assay were then normalized to cell number. ATP production rates (J_ATP_) were calculated based on protocols developed by Mookerjee *et al*., and adapted by Ma *et al* ^40,41^. J_ATP_ by glycolysis (J_ATP_glyc) and oxidative metabolism (J_ATP_ox) were calculated by ECAR or OCR, respectively, before any drug treatment for basal metabolism, and after FCCP treatment for maximal respiration. J_ATP_total represent the sum of J_ATP_glyc and J_ATP_ox.

### Stable isotope labeling (SIL) and metabolomics

DCs were cultured for the indicated times in medium containing 10% dialyzed FBS, 2 mM L-glutamine, 100 U/mL penicillin/streptomycin, 55 μM β-mercaptoethanol, and 10 mM ^13^C-labeled glucose (Cambridge Isotope Laboratories). Cells were washed with ice-cold saline and snap frozen on dry-ice and stored at −80°C.

Metabolites were extracted by modified Bligh-Dyer extraction ^42^ by the addition of ice-cold methanol (A456, Fisher Scientific) directly to frozen cells, to which one volume of chloroform (A456, Fisher Scientific) was added. The sample was vortexed for 10 s, incubated on ice for 30 min, and then 0.9 parts of LC/MS grade water (W6-4, Thermo Fisher Scientific) was added. The samples were vortexed vigorously and centrifuged at maximum speed to achieve phase separation. The top layer containing polar metabolites was aliquoted into a fresh tube and dried in a speedvac for LC/MS analysis. The bottom layer was retained for fatty acid methyl-ester measurement.

For LC-MS analysis, metabolite extracts were resuspended in 50 μL of 60% acetonitrile (A955, Fisher Scientific) and analyzed by high resolution accurate mass spectrometry using an ID-X Orbitrap mass spectrometer (Thermo Fisher Scientific) coupled to a Thermo Vanquish Horizon liquid chromatography system. 2 μL of sample volume was injected on column. Chromatographic separations were accomplished with Acquity BEH Amide (1.7 μm, 2.1 mm x 150 mm) analytical columns (#176001909, Waters, Eschborn, Germany) fitted with a pre-guard column (1.7 μm, 2.1mm x 5 mm; #186004799, Waters) using an elution gradient with a binary solvent system. Solvent A consisted of LC/MS grade water (W6-4, Fisher), and Solvent B was 90% LC/MS grade acetonitrile (A955, Fisher). For negative mode analysis, both mobile phases contained 10 mM ammonium acetate (A11450, Fisher Scientific), 0.1% (v/v) ammonium hydroxide, and 5 μM medronic acid (5191-4506, Agilent Technologies). For positive mode analysis, both mobile phases contained 10 mM ammonium formate (A11550, Fisher), and 0.1% (v/v) formic acid (A11710X1, Fisher). For both negative and positive mode analyses the 20-min analytical gradient at a flow rate of 400 μL/min was: 0–1.0 min ramp from 100% B to 90% B, 1.0–12.5 min from 90% B to 75% B, 12.5–19 min from 75% B to 60% B, and 19–20 min hold at 60% B. Following every analytical separation, the column was re-equilibrated for 20 min as follows: 0– 1 min hold at 65%B at 400 μL/min, 1–3 min hold at 65% B and ramp from 400 μL/min to 800 μL/min, 3–14 min hold at 65% B and 800 μL/min, 14–14.5 min ramp from 65% B to 100% B at 800 μL/min, 14.5–16 min hold at 100% B and increase flow from 800 μL/min to 1200 μL/min, 16–18.4 min hold at 100% B at 1200 μL/min, 18.4–19.5 min hold at 100% B and decrease flow from 1200 μL to 400 μL/min, 19.5–20 min hold at 100% B and 400 μL/min. The column temperature was maintained at 40°C. The H-ESI source was operated at spray voltage of 2500 V(negative mode)/3500 V(positive mode), sheath gas: 60 a.u., aux gas: 19 a.u., sweep gas: 1 a.u., ion transfer tube: 300°C, vaporizer: 300°C. For isotopically labelled experimental replicates, high resolution MS^1^ data was collected with a 20-min full-scan method with m/z scan range using quadrupole isolation from 70 to 1000, mass resolution of 120,000 FWHM, RF lens at 35%, and standard automatic gain control (AGC). Unlabelled control samples were used for data dependent MS^2^ (ddMS^2^) fragmentation for compound identification and annotation via the AquireX workflow (Thermo Scientific). In this workflow, first blank and experimental samples are injected to generate exclusion and inclusion lists, respectively, followed by iterative sample injections for ddMS^2^ fragmentation where triggered ions are added to the exclusion list for subsequent injections. ddMS^2^ data was collected using MS1 resolution at 60,000, MS^2^ resolution at 30,000, intensity threshold at 2.0 x 10^4^, and dynamic exclusion after one trigger for 10 s. MS^2^ fragmentation was completed first with HCD using stepped collision energies at 20, 35, and 50% and was followed on the next scan by CID fragmentation in assisted collision energy mode at 15, 30, and 45% with an activation Q of 0.25. Both MS^2^ scans used standard AGC and a maximum injection time of 54ms. The total cycle time of the MS^1^ and ddMS^2^ scans was 0.6 s.

Full scan LC-MS data were analyzed in Compound Discoverer (v 3.2, Thermo Scientific). Compounds were identified by chromatography specific retention time of external standards and MS^2^ spectral matching using the mzCloud database (Thermo Scientific).

TCA cycle intermediates were measured via GC-MS following LC-MS analysis. Briefly, following LC/MS extracts were dried and derivatized with 30 μL of methoxyamine (11.4 mg/mL) in pyridine and 70 μL of MTBSFA+1%TMCS as described previously ^43^. In addition, GC-MS was used to evaluate incorporation of 13C into de novo synthesized fatty acids from metabolic precursors using fatty acid methyl-esters (FAMEs). The bottom organic fraction from the Bligh-Dyer extraction (above) was aliquoted, dried in a speedvac, and FAMEs were generated as described previously. GC-MS analysis of both TBDMS derivatives and FAMEs were conducted on an Agilent 7890/5977b GC/MSD equipped with a DB-5MS+DG (30 m x 250 μm x 0.25 μm) capillary column (Agilent J&W, Santa Clara, CA, USA) was used. Data were collected by electron impact set at 70 eV. A total of 1 μL of the derivatized sample was injected in the GC in split mode (1:2 or 1:4) with inlet temperature set to 280°C, using helium as a carrier gas with a column flow rate of 1.2 mL/min. The oven program for all metabolite analyses started at 95°C for 1 min, increased at a rate of 40°C/min until 118°C and held for 2 min, then increased to 250°C at a rate of 12°C/min, then increased to 320°C at a rate of 40°C/min and finally held at 320°C for 7 min. The source temperature was 230°C, the quadrupole temperature was 150°C, and the GC/MS interface at 285°C. Data were acquired both in scan mode (50–800 m/z) and 2 Hz.

MassHunter software (v10, Agilent Technologies) was used for peak picking and integration of GC/MS data. Peak areas of all isotopologues for a molecular ion of each compound in both labeled experimental and unlabeled control samples were used for mass isotopologue distribution analysis via a custom algorithm developed at VAI. This algorithm uses matrices correcting for natural contribution of isotopologue enrichment were generated for each metabolite as described previously ^45,46^.

### RNA sequencing and analysis

RNA was isolated using TRIzol (ThermoFisher Scientific) according to manufacturer’s instructions. Libraries were prepared by the Van Andel Genomics Core from 500 ng of total RNA using the KAPA RNA HyperPrep Kit with RiboseErase (v2.17) (Kapa Biosystems, Wilmington, MA USA). RNA was sheared to 300-400 bp. Prior to PCR amplification, cDNA fragments were ligated to IDT for Illumina DNA/RNA UD Indexed adapters (Illumina Inc, San Diego CA, USA). Quality and quantity of the finished libraries were assessed using a combination of Agilent DNA High Sensitivity chip (Agilent Technologies, Inc.) and QuantiFluor^®^ dsDNA System (Promega Corp., Madison, WI, USA. Individually indexed libraries were pooled and 100 bp, paired end sequencing was performed on an Illumina NovaSeq6000 sequencer using an S1, 200 bp sequencing kit (Illumina Inc., San Diego, CA, USA) to an average depth of 50M reads per sample. Base calling was done by Illumina RTA3 and demultiplexed to FASTQ format with Illumina Bcl2fastq v1.9.0

Raw reads were first trimmed of adapters with trim_galore (https://www.bioinformatics.babraham.ac.uk/projects/trim_galore/) and then mapped to mm10 with STAR v2.7.3a using option –quantMode GeneCounts to directly output counts for all features from GENCODE release M20^47^. A negative binomial generalized log-linear model was then fit to the filtered count data with edgeR using the weighted trimmed mean of M-values to normalize for library size and composition biases ^48^. An interactions model was fit to find genes that responded differently depending on the stimulation type ((treatmentIFNb-treatmentcontrol)-(treatmentLPS-treatmentcontrol)). P-values were generated using empirical Bayes quasi-likelihood F-tests, and then adjusted using the BH method; adjusted P-values less than 0.05 were considered significant.

Heatmaps were generated from library-size normalized counts centered across genes (z-scores). Heatmaps were generated using the R package ComplexHeatmap ^49^. KEGG and REACTOME pathways were retrieved with the R package msigdbr^50^. Pathway enrichment testing was done with GSEA as implemented in the clusterProfiler v3.14.3 function ‘GSEA.’ Enrichment testing for GO terms was also done with clusterProfiler v3.14.3 using the enrichGO function.

### Quantitative RT-PCR

RNA was isolated using TRIzol (ThermoFisher Scientific) according to manufacturer’s instructions. cDNA was obtained using RT MasterMix (Applied Biological Materials) and used to perform a SYBR-based real-time PCR (BioLine) with primers (Table 1) from Integrated DNA Technologies. Data were generated using the ΔΔCq method. Relative gene expression was normalized to that of hypoxanthine-guanine phosphoribosyltransferase (HPRT).

**Table 1.**
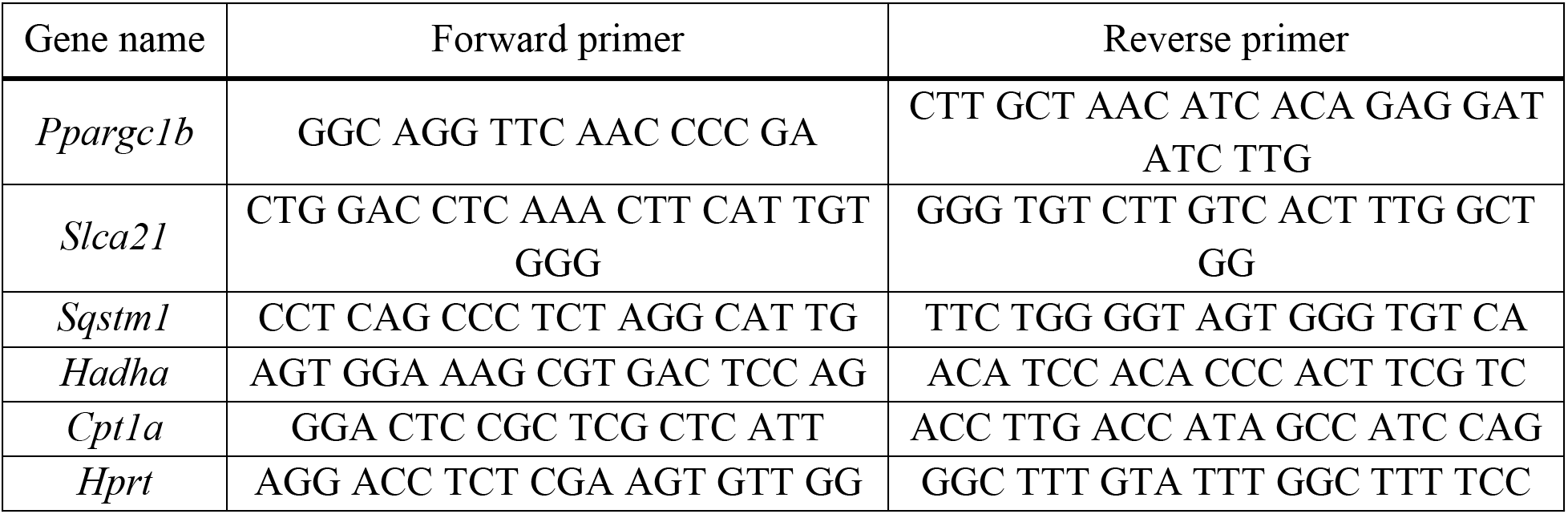
List of primers used for qPCR

### Flow cytometry

Cells were washed with wash buffer (PBS containing 1% FBS, 1 mM EDTA, and 0.05% sodium azide) and stained with eFluor 506 Fixable Viability Dye (Invitrogen) and Fc block (anti-CD16/CD32) prior to staining with cell surface markers, which include human CD8 (OKT-8), CD11c (N418), MHC-II (M5/144.15.2), CD86 (GL-1), and PD-L1 (10F.9G2). To examine intracellular cytokine IL-12p40 (C17.8), cells were fixed with IC Fixation Buffer (Invitrogen) for 30 minutes, followed by cell permeabilization using Permeabilization Buffer (Invitrogen), and at least 1 hour of intracellular cytokine staining in Permeabilization Buffer. For mitochondrial mass measurements, cells were stained with 50 nM MitoSpy Green (BioLegend) or 50 nM MitoTracker Deep Red (ThermoFisher Scientific) in HBSS at 37°C for 20 minutes, followed by viability and surface marker staining as described above. Apotracker Green (BioLegend) was used at 400 nM according to manufacturer’s instructions to detect apoptotic cells. Samples were acquired in wash buffer on the Cytoflex (Beckman Coulter) or Aurora (Cytek) and analyzed using FlowJo software.

### Western blot

Protein lysates were prepared in RIPA lysis buffer with protease inhibitor and phosphatase inhibitor, then quantified by detergent compatible protein assay (Bio-Rad). Samples were run on 10% or 12.5% gels using SDS-polyacrylamide gel electrophoresis, and proteins were transferred by Turbo transfer system (Bio-Rad) onto methanol-activated polyvinylidine fluoride membranes. After blocking for 1 hour with 5% milk, membranes were incubated overnight rocking at 4°C with antibodies against β-actin (13E5), LC3I/II, SQSTM1 (D1Q5S), and BNIP3 (all from Cell Signaling Technology), and Total OXPHOS Rodent WB Antibody Cocktail (both from Abcam), all at 1:1000 in 4% BSA. Following washes with Tris-buffered saline (TBS) containing 0.1% Tween-20 (TBS-T), membranes were incubated with horseradish peroxidase-linked antibody to rabbit or mouse IgG in 5% milk for 45 minutes. Enhanced chemiluminescence using SuperSignal substrate (ThermoFisher Scientific) and ChemiDoc (Bio-Rad) were used to develop blots. Original blots are presented in Supplementary Figures.

### Statistical analysis

Data were analyzed using GraphPad Prism software (version 9). An unpaired or paired Student’s *t*-test was performed as indicated to determine statistical significance between two conditions. A two-way analysis of variance was performed to determine statistical significance between multiple groups. Differences between conditions were considered significant when *P* values were below 0.05.

## Author contributions

H.G., J.S-P., R.G.J, and C.M.K. conceived this study. H.G., R.S., E.H.M, and C.M.K designed the experiments. H.G. and A.V. performed experiments. H.G., R.S., I.B., E.H.M and H.S. analyzed data and generated figures. H.G. wrote the manuscript. H.G., J.S-P., and C.M.K. edited the manuscript.

## Acknowledgements

The authors thank current and former members of the C.M.K and R.G.J. laboratories for thoughtful discussion. We also thank Dr. Sara Nowinski for helpful feedback of the manuscript. The authors acknowledge the staff and services of the Flow Cytometry, Genomics, Metabolomics and Bioenergetics, and Vivarium cores at the Van Andel Research Institute. We also acknowledge Alicia Withrow at the Center for Advanced Microscopy at Michigan State University for the mitochondrial images. This study was supported in part by the Canadian Institutes of Health Research grant to C.M.K. (MOP-126184).

## Competing interests

The authors have no conflicts of interest to declare.

## Data availability

Data will be available upon request. Submission numbers for RNAseq will be supplied before publication.

## Notes

### Competing Interest Statement

The authors have declared no competing interest.

## References

1. Krawczyk, C. M. et al. Toll-like receptor–induced changes in glycolytic metabolism regulate dendritic cell activation. Blood 115, 4742–4749 (2010).

2. O’Neill, L. A. J., Kishton, R. J. & Rathmell, J. A guide to immunometabolism for immunologists. Nat. Rev. Immunol. 16, 553–565 (2016).

3. Buck, M. D., Sowell, R. T., Kaech, S. M. & Pearce, E. L. Metabolic Instruction of Immunity. Cell 169, 570–586 (2017).

4. Lin, J., Handschin, C. & Spiegelman, B. M. Metabolic control through the PGC-1 family of transcription coactivators. Cell Metab. 1, 361–370 (2005).

5. Rhee, J. et al. Regulation of hepatic fasting response by PPARγ coactivator-1α (PGC-1): Requirement for hepatocyte nuclear factor 4α in gluconeogenesis. Proc. Natl. Acad. Sci. U. S. A. 100, 4012–4017 (2003).

6. Lin, J. et al. Hyperlipidemic Effects of Dietary Saturated Fats Mediated through PGC-1β Coactivation of SREBP. Cell 120, 261–273 (2005).

7. St-Pierre, J. et al. Bioenergetic Analysis of Peroxisome Proliferator-activated Receptor γ Coactivators 1α and 1β (PGC-1α and PGC-1β) in Muscle Cells. J. Biol. Chem. 278, 26597–26603 (2003).

8. Dumauthioz, N. et al. Enforced PGC-1α expression promotes CD8 T cell fitness, memory formation and antitumor immunity. Cell. Mol. Immunol. 1–11 (2020) doi:10.1038/s41423-020-0365-3.

9. Scharping, N. E. et al. The Tumor Microenvironment Represses T Cell Mitochondrial Biogenesis to Drive Intratumoral T Cell Metabolic Insufficiency and Dysfunction. Immunity 45, 374–388 (2016).

10. Bengsch, B. et al. Bioenergetic Insufficiencies Due to Metabolic Alterations Regulated by the Inhibitory Receptor PD-1 Are an Early Driver of CD8+ T Cell Exhaustion. Immunity 45, 358–373 (2016).

11. Vats, D. et al. Oxidative metabolism and PGC-1β attenuate macrophage-mediated inflammation. Cell Metab. 4, 13–24 (2006).

12. Everts, B. et al. TLR-driven early glycolytic reprogramming via the kinases TBK1-IKKε supports the anabolic demands of dendritic cell activation. Nat. Immunol. 15, 323–332 (2014).

13. Zhao, F. et al. Paracrine Wnt5a-β-Catenin Signaling Triggers a Metabolic Program that Drives Dendritic Cell Tolerization. Immunity 48, 147–160.e7 (2018).

14. Malinarich, F. et al. High Mitochondrial Respiration and Glycolytic Capacity Represent a Metabolic Phenotype of Human Tolerogenic Dendritic Cells. J. Immunol. 194, 5174–5186 (2015).

15. Pantel, A. et al. Direct Type I IFN but Not MDA5/TLR3 Activation of Dendritic Cells Is Required for Maturation and Metabolic Shift to Glycolysis after Poly IC Stimulation. PLOS Biol. 12, e1001759 (2014).

16. Everts, B. et al. Commitment to glycolysis sustains survival of NO-producing inflammatory dendritic cells. Blood 120, 1422 (2012).

17. Ahmed, D. et al. Transcriptional Profiling Suggests Extensive Metabolic Rewiring of Human and Mouse Macrophages during Early Interferon Alpha Responses. Mediators Inflamm. 2018, e5906819 (2018).

18. Mansouri, S. et al. In vivo reprogramming of pathogenic lung TNFR2+ cDC2s by IFNβ inhibits HDM-induced asthma. Sci. Immunol. 6, eabi8472 (2021).

19. Guak, H. et al. Glycolytic metabolism is essential for CCR7 oligomerization and dendritic cell migration. Nat. Commun. 9, 2463 (2018).

20. Lin, S.-C. & Hardie, D. G. AMPK: Sensing Glucose as well as Cellular Energy Status. Cell Metab. 27, 299–313 (2018).

21. Jäger, S., Handschin, C., St.-Pierre, J. & Spiegelman, B. M. AMP-activated protein kinase (AMPK) action in skeletal muscle via direct phosphorylation of PGC-1α. Proc. Natl. Acad. Sci. U. S. A. 104, 12017–12022 (2007).

22. Hardie, D. G. AMP-activated/SNF1 protein kinases: conserved guardians of cellular energy. Nat. Rev. Mol. Cell Biol. 8, 774–785 (2007).

23. Jornayvaz, F. R. & Shulman, G. I. Regulation of mitochondrial biogenesis. Essays Biochem. 47, 69–84 (2010).

24. Wolfrum, C. & Stoffel, M. Coactivation of Foxa2 through Pgc-1β promotes liver fatty acid oxidation and triglyceride/VLDL secretion. Cell Metab. 3, 99–110 (2006).

25. An, J. et al. CHCM1/CHCHD6, Novel Mitochondrial Protein Linked to Regulation of Mitofilin and Mitochondrial Cristae Morphology *. J. Biol. Chem. 287, 7411–7426 (2012).

26. Russell, A. P., Hesselink, M. K. C., Lo, S. K. & Schrauwen, P. Regulation of metabolic transcriptional co-activators and transcription factors with acute exercise. FASEB J. 19, 986–988 (2005).

27. Rowe, G. C. et al. Disconnecting mitochondrial content from respiratory chain capacity in PGC-1 deficient skeletal muscle. Cell Rep. 3, 1449–1456 (2013).

28. Eisele, P. S., Salatino, S., Sobek, J., Hottiger, M. O. & Handschin, C. The Peroxisome Proliferator-activated Receptor γ Coactivator 1α/β (PGC-1) Coactivators Repress the Transcriptional Activity of NF-κB in Skeletal Muscle Cells. J. Biol. Chem. 288, 2246–2260 (2013).

29. Eisele, P. S., Furrer, R., Beer, M. & Handschin, C. The PGC-1 coactivators promote an anti-inflammatory environment in skeletal muscle in vivo. Biochem. Biophys. Res. Commun. 464, 692–697 (2015).

30. Chang, W.-C. et al. Genetic variants of PPAR-gamma coactivator 1B augment NLRP3-mediated inflammation in gouty arthritis. Rheumatology 56, 457–466 (2017).

31. Salazar, G. et al. SQSTM1/p62 and PPARGC1A/PGC-1alpha at the interface of autophagy and vascular senescence. Autophagy 16, 1092–1110.

32. Sopariwala, D. H. et al. Long-term PGC1β overexpression leads to apoptosis, autophagy and muscle wasting. Sci. Rep. 7, 10237 (2017).

33. Vainshtein, A., Tryon, L. D., Pauly, M. & Hood, D. A. Role of PGC-1α during acute exercise-induced autophagy and mitophagy in skeletal muscle. Am. J. Physiol.-Cell Physiol. 308, C710–C719 (2015).

34. Mills, E. L. et al. Succinate Dehydrogenase Supports Metabolic Repurposing of Mitochondria to Drive Inflammatory Macrophages. Cell 167, 457–470.e13 (2016).

35. Bailis, W. et al. Distinct modes of mitochondrial metabolism uncouple T cell differentiation and function. Nature 571, 403–407 (2019).

36. Shin, B. et al. Mitochondrial Oxidative Phosphorylation Regulates the Fate Decision between Pathogenic Th17 and Regulatory T Cells. Cell Rep. 30, 1898–1909.e4 (2020).

37. Fernandez-Vizarra, E. & Zeviani, M. Mitochondrial disorders of the OXPHOS system. FEBS Lett. 595, 1062–1106 (2021).

38. Letts, J. A. & Sazanov, L. A. Clarifying the supercomplex: the higher-order organization of the mitochondrial electron transport chain. Nat. Struct. Mol. Biol. 24, 800–808 (2017).

39. Paddison, P. J. et al. Cloning of short hairpin RNAs for gene knockdown in mammalian cells. Nat. Methods 1, 163–167 (2004).

40. Mookerjee, S. A., Gerencser, A. A., Nicholls, D. G. & Brand, M. D. Quantifying intracellular rates of glycolytic and oxidative ATP production and consumption using extracellular flux measurements. J. Biol. Chem. 292, 7189–7207 (2017).

41. Ma, E. H. et al. Metabolic Profiling Using Stable Isotope Tracing Reveals Distinct Patterns of Glucose Utilization by Physiologically Activated CD8+ T Cells. Immunity 51, 856–870.e5 (2019).

42. Bligh, E. G. & Dyer, W. J. A rapid method of total lipid extraction and purification. Can. J. Biochem. Physiol. 37, 911–7 (1959).

43. Ma, E. H. et al. Metabolic Profiling Using Stable Isotope Tracing Reveals Distinct Patterns of Glucose Utilization by Physiologically Activated CD8+ T Cells. Immunity 51, 856–870.e5 (2019).

44. Griss, T. et al. Metformin Antagonizes Cancer Cell Proliferation by Suppressing Mitochondrial-Dependent Biosynthesis. PLoS Biol 13, e1002309 (2015).

45. Trefely, S., Ashwell, P. & Snyder, N. W. FluxFix: automatic isotopologue normalization for metabolic tracer analysis. BMC Bioinformatics 17, 485 (2016).

46. Fernandez, C. A., Des Rosiers, C., Previs, S. F., David, F. & Brunengraber, H. Correction of 13C mass isotopomer distributions for natural stable isotope abundance. J. Mass Spectrom. JMS 31, 255–62 (1996).

47. Dobin, A. et al. STAR: ultrafast universal RNA-seq aligner. Bioinforma. Oxf. Engl. 29, 15–21 (2013).

48. Robinson, M. D., McCarthy, D. J. & Smyth, G. K. edgeR: a Bioconductor package for differential expression analysis of digital gene expression data. Bioinformatics 26, 139–140 (2010).

49. Gu, Z., Eils, R. & Schlesner, M. Complex heatmaps reveal patterns and correlations in multidimensional genomic data. Bioinformatics 32, 2847–2849 (2016).

50. Dolgalev, I. msigdbr: MSigDB Gene Sets for Multiple Organisms in a Tidy Data Format. (2021).

