## Supplemental Data for "PGC-1β maintains mitochondrial metabolism and restrains inflammatory gene expression"

Supplementary Figures

Figure S1

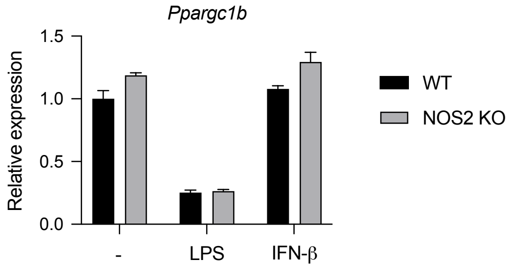


Figure S1 – NOS2 expression does not affect *Ppargc1b* expression

DCs derived from wild-type (WT) or *Nos2-/-* (KO) mice were left unstimulated or stimulated with LPS (100 ng/mL) or IFN-β (1000 U/mL) for 18 hours. Fold changes of *Ppargc1b* expression were made relative to *Hprt*. Data are from one experiment representative of two experiments.

Figure S2

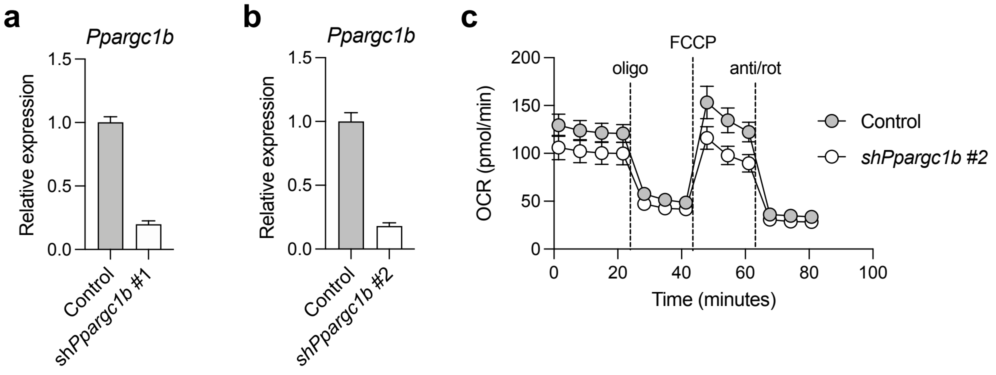


Figure S2 – Confirmation of knockdown by *Ppargc1b* shRNA

DCs were transduced with control shRNA or two different *Ppargc1b* shRNA (hairpin #1 in **(a)** and hairpin #2 in **(b)**), and knockdown was confirmed by RT-qPCR. **(c)** OCR profile of DCs transduced with control shRNA or *Ppargc1b* hairpin #2 with sequential treatments of oligomycin (oligo), FCCP, and antimycin/rotenone (anti/rot). Data are of one experiment representative of **(a)** > three experiments or **(b,c)** two experiments.

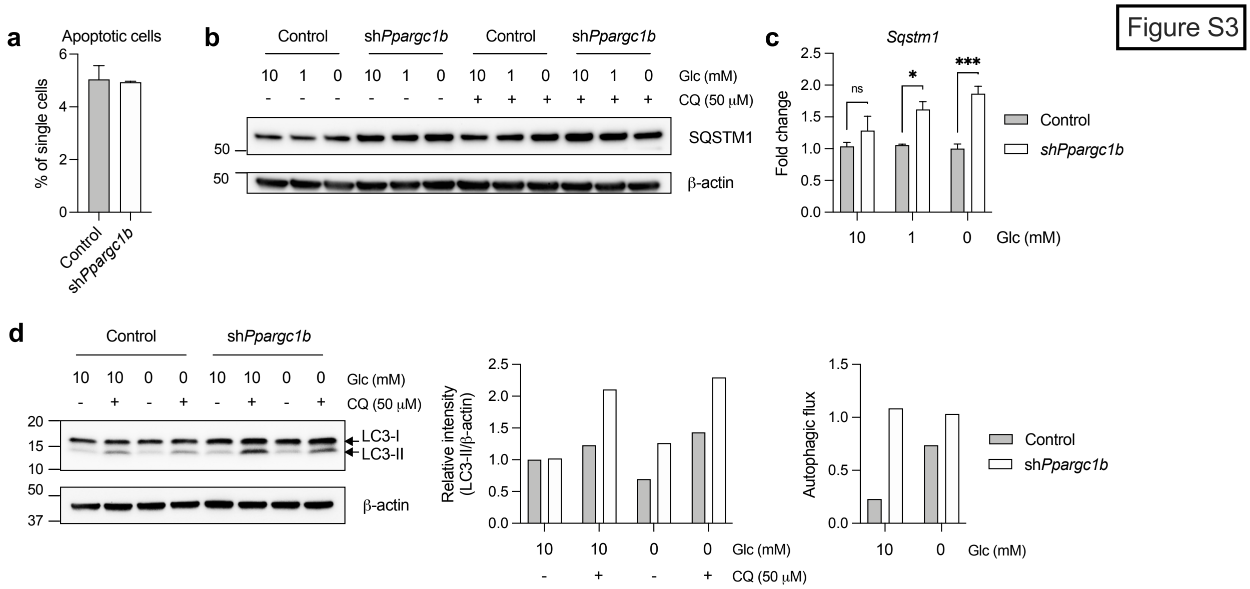


Figure S3 – PGC-1β deficiency increases autophagy and does not affect apoptosis

DCs were transduced with control or *Ppargc1b* shRNA and **(a)** examined for apoptosis, represented by the frequency of cells that were positive for both ApoTracker Green and eFluor 506 Fixable Viability Dye. DCs were cultured in 0, 1 or 10 mM glucose as indicated for 6 hours, with 50 μM chloroquine (CQ) added for the last 2 hours of culture, and **(b)** SQSTM1 was visualized by western blot. **(c)** Gene expression of *Sqstm1* relative to *Hprt*. **(d)** LC3-I/II visualized by western blot (left) and ratio of relative intensity of LC3-II to β-actin (middle). Autophagic flux was calculated by determining the difference between the intensity of LC3-II normalized by β-actin in the presence and absence of chloroquine (right). Data are of one experiment representative of **(a)** one experiment (mean and s.d. of triplicates), **(b-d)** three experiments. Statistical significance in **(c)** was determined by two-way ANOVA. * $p < 0.05$, *** $p > 0.001$

Figure S4

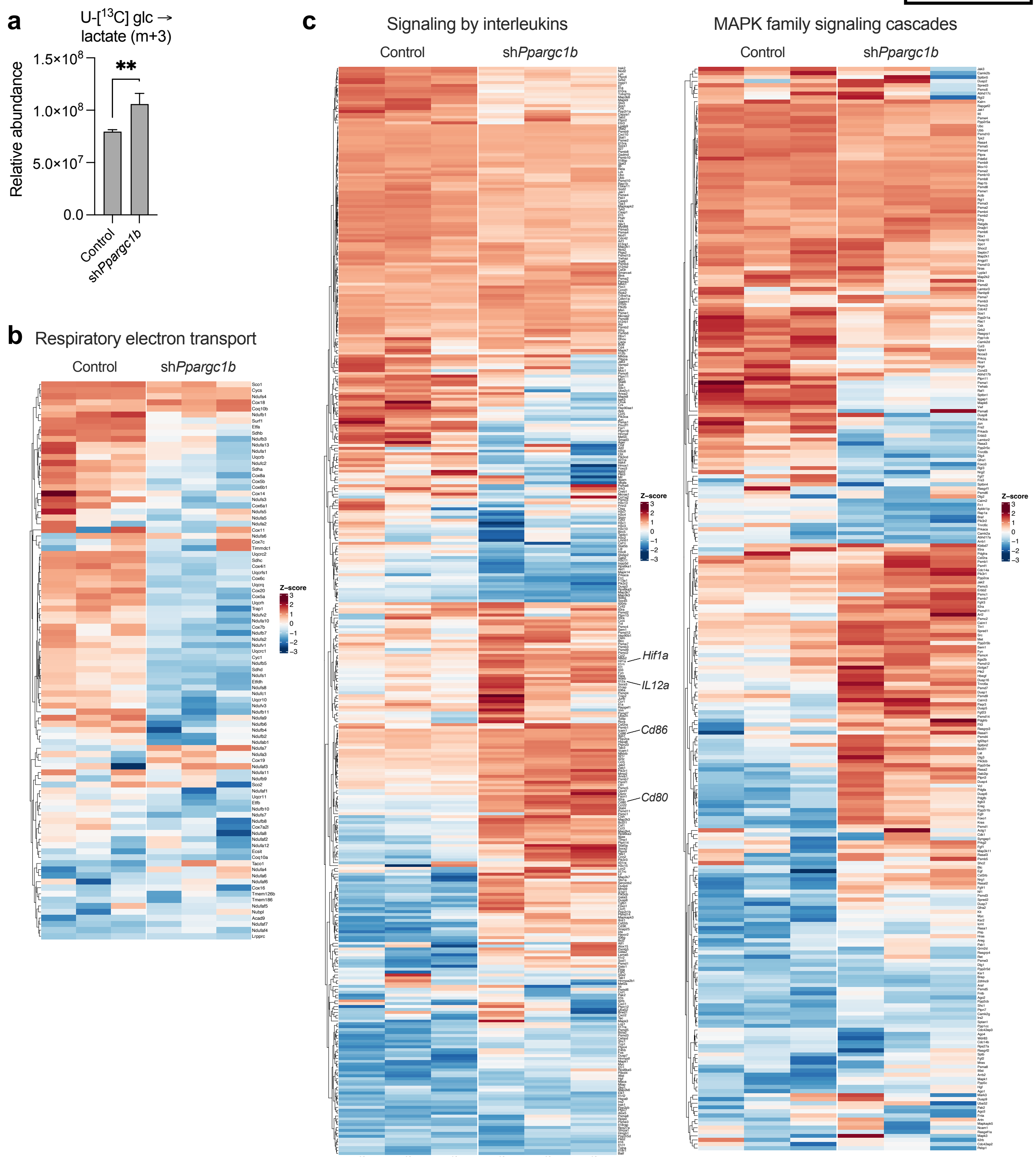


<u>Figure S4 – Metabolite analysis following $^{13}$C-glucose pulse and gene expression analysis of PGC-1β-deficient DCs</u>

**(a)** Relative abundance of labeled ($^{13}$C) intracellular lactate (m+3 isotopomer) from DCs transduced with control or *Ppargc1b* shRNA and cultured with U-[$^{13}$C]-glucose for 6 h prior to metabolite extraction. **(b,c)** Heat maps showing the expression of genes involved in **(b)** respiratory electron transport and **(c)** signaling by interleukins (left) and MAPK family signaling cascades (right). Data in **(a)** are of one experiment representative of two experiments (mean and s.e.m. of triplicates). ** $p > 0.01$

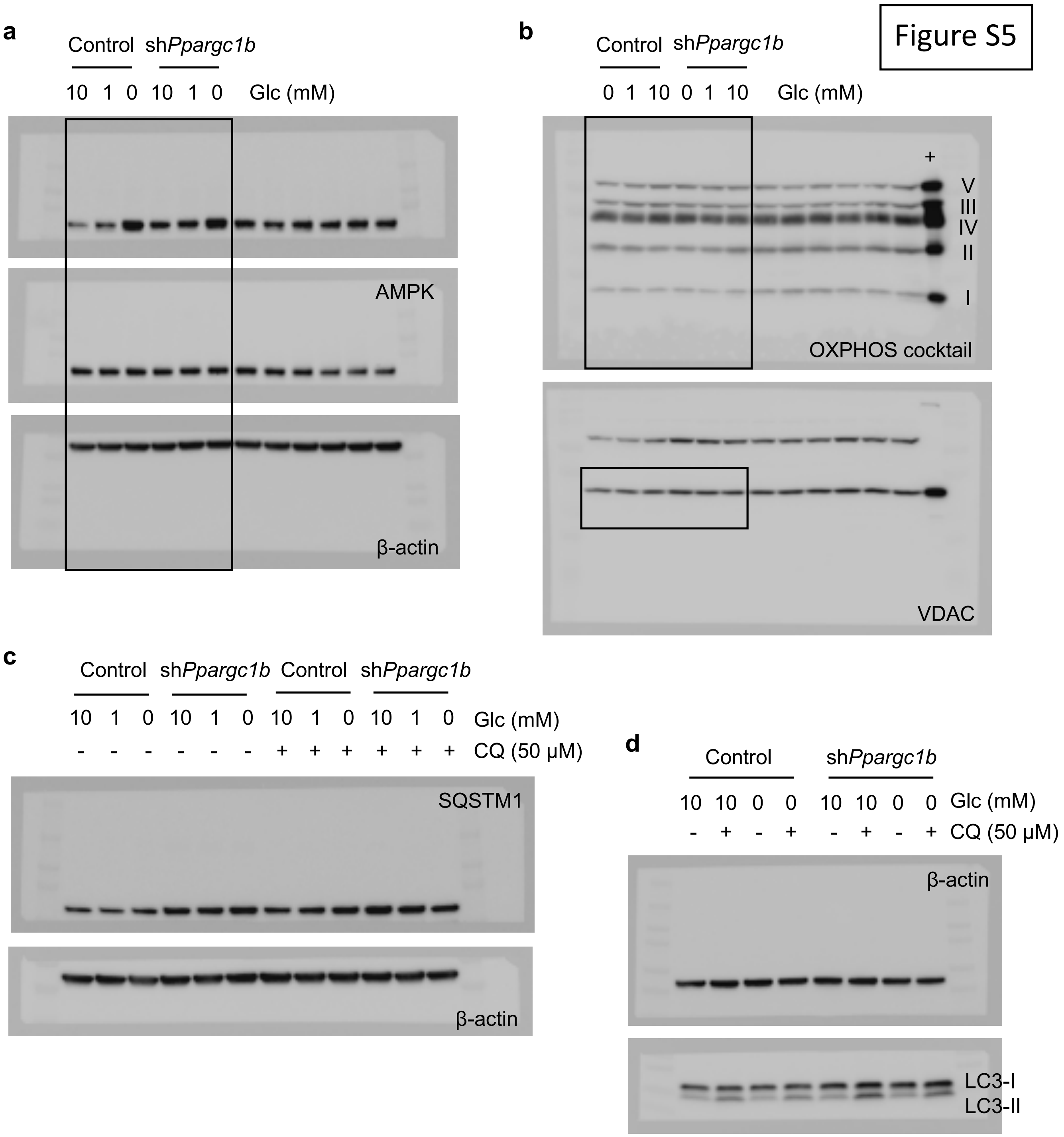


Figure S5 – Uncropped blots

Uncropped blots of **(a)** p-AMPK, AMPK, and β-actin corresponding to Fig. 3g, **(b)** OXPHOS cocktail (ETC complexes I-V) and VDAC corresponding to Fig. 4b, **(c)** SQSTM1 and β-actin corresponding to Supplementary Fig. S3b, and **(d)** β-actin and LC3-I/II corresponding to Supplementary Fig. S3d.
